# Cytoplasmic TDP-43 leads to early functional impairments without neurodegeneration in a Serotonergic Neuron-Specific *C. elegans* Model

**DOI:** 10.1101/2025.07.30.667669

**Authors:** Ailín Lacour, Florencia Vassallu, Diego Rayes, Lionel Muller Igaz

## Abstract

TDP-43 proteinopathies, such as amyotrophic lateral sclerosis (ALS) and frontotemporal dementia (FTD), are marked by the pathological cytoplasmic accumulation of TAR DNA-binding protein 43 (TDP-43), leading to progressive neuronal dysfunction and degeneration. To investigate the early functional consequences of TDP-43 mislocalization, we generated *Caenorhabditis elegans* models expressing either wild-type human TDP-43 or a variant with a mutated nuclear localization signal (ΔNLS), specifically in serotonergic neurons. These neurons were chosen because i) serotonin deficits are a feature of ALS/FTD and ii) in *C. elegans*, they regulate well-characterized behaviors, providing a straightforward readout of neuronal function. We found that expression of either TDP-43 variant impaired serotonin-dependent behaviors—including pharyngeal pumping, egg-laying, and locomotion slowing upon food encounter—with the cytoplasmic ΔNLS form causing more severe deficits. Serotonergic neurons remained i) morphologically intact, indicating that neuronal dysfunction precedes overt neurodegeneration; and ii) partially responsive to the selective serotonin reuptake inhibitor fluoxetine, suggesting that neurotransmitter release is still partially functional. Altogether, our findings demonstrate that cytoplasmic TDP-43 disrupts neuronal signaling and behavior early in disease progression. This *C. elegans* model provides a genetically tractable system to dissect early mechanisms of TDP-43-mediated dysfunction and to identify therapeutic strategies targeting predegenerative stages of ALS/FTD.

## Introduction

TAR DNA-binding protein 43 (TDP-43) is an evolutionarily conserved, ubiquitously expressed RNA-binding protein that plays critical roles in RNA metabolism, including transcription, splicing, transport, and stability [1]. Under normal physiological conditions, TDP-43 is predominantly localized in the nucleus. However, in neurodegenerative disorders such as amyotrophic lateral sclerosis (ALS) and frontotemporal dementia (FTD), TDP-43 undergoes pathological mislocalization to the cytoplasm, where it forms insoluble aggregates - a hallmark feature of these pathological conditions [2, 3].

While ALS has been classically characterized by the degeneration of motor neurons [4], emerging evidence suggests that other neuronal populations, including serotonergic neurons, may also be affected [5]. Studies in ALS mouse models have reported cell-autonomous pathological alterations in serotonergic neurons, and pharmacological modulation of serotonin signaling has demonstrated partial therapeutic benefits [6, 7]. These observations indicate that non-motor neuronal systems may contribute to disease pathogenesis, although their precise role remains unclear.

The nematode *Caenorhabditis elegans* has emerged as a powerful model organism for studying neurodegenerative processes, offering several unique advantages: a fully mapped nervous system consisting of exactly 302 neurons in the hermaphrodite, genetic tractability, and a short lifespan [8, 9]. Importantly, *C. elegans* exhibits well-characterized behaviors that are controlled by specific, identifiable neuronal circuits, allowing for precise assessment of neuronal function through behavioral assays [10-12]

Previous studies have successfully modeled various proteinopathies in *C. elegans*, including Parkinson’s disease, Alzheimer’s disease, and TDP-43 proteinopathies [13, 14]. These models commonly feature pan-neuronal or muscle-specific expression of human proteins (α-synuclein, β-amyloid, and TDP-43) linked to these pathologies and frequently use locomotion as the primary behavioral readout [15, 16]. While valuable, such approaches may not capture more subtle aspects of neuronal dysfunction.

In this study, we focused on the serotonergic system of *C. elegans*, which consists of just three pairs of well-defined neurons that control quantifiable behaviors including feeding (pharyngeal pumping), locomotion modulation, and egg-laying [17-22]. The simplicity and well-characterized nature of this system make it particularly suitable for investigating how TDP-43 pathology affects specific neuronal circuits.

We generated transgenic *C. elegans* strains expressing either human wild-type TDP-43 (*hTDP-43-WT*) or a nuclear exclusion mutant (*hTDP-43-ΔNLS*, which lacks a functional nuclear localization signal and is retained in the cytoplasm) [23] specifically in serotonergic neurons. This targeted approach allowed us to address two key questions: (1) How does cytoplasmic accumulation of TDP-43 affect the function of a defined neuronal population? (2) Are behavioral deficits associated with structural neurodegeneration or more subtle functional impairments?

Our results demonstrate that cytoplasmic accumulation of TDP-43 disrupts serotonergic neuron function, leading to measurable behavioral impairments while preserving neuronal integrity. These findings not only establish a novel animal model for studying TDP-43 proteinopathies but also provide insights into the mechanisms by which TDP-43 dysregulation affects neuronal function in specific circuits.

## Materials and methods

### *C. elegans* culture and maintenance

All *C. elegans* strains were grown at room temperature (22ºC) on nematode growth media (NGM) agar plates with *Escherichia coli* OP50 as a food source. The wild-type reference strain used in this study is N2 Bristol. Some of the strains were obtained through the Caenorhabditis Genetics Center (CGC, University of Minnesota). Worm population density was maintained at a low level throughout their development and during the assays. All experiments were conducted on age-synchronized animals. This was achieved by placing gravid worms on NGM plates and removing them after two hours. The assays were performed on the animals hatched from the eggs laid in these two hours.

Transgenic strains were generated by microinjection of plasmid DNA containing the construct *Ptph-1::hTDP43* and cDNA *Ptph-1::hTDP43(ΔNLS)* at 10 ng/µL into the germ line of *lin-15 (n765ts*) mutants with the co-injection marker *lin-15* rescuing plasmid pL15EK (80 ng/µl). At least three independent transgenic lines were obtained. Data are shown from a single representative line.

The strains used were:

N2 (wild-type)

MT15434 *tph-1(mg280) II*,

MT13471 *[ptph-1::GFP]*

MT1082 *egl-1(n487) V*.

OAR165 *lin-15(n765ts); nbaEx20[Ptph-1::hTDP43(10) +lin-15 (80)*]

OAR166 *lin-15(n765ts); nbaEx21[Ptph-1::hTDP43(ΔNLS)) +lin-15 (80)]*

OAR199 *lin-15(n765ts); nbaEx20[Ptph-1::hTDP43(10) +lin-15 (80)]; ptph-1::GFP*

OAR200 *lin-15(n765ts); nbaEx21[Ptph-1::hTDP43(ΔNLS)) +lin-15 (80)]; ptph-1::GFP*

### Microscopy and image analysis

For microscopy, L4 age-synchronized animals were mounted in M9 with levamisole (10 mM) onto slides with 2% agarose pads. Images were acquired on confocal microscopy (LSCM; Leica DMIRE2) with 20X and 63X objectives. For *tph-1*::GFP expression levels analysis, animals containing the corresponding transcriptional GFP reporter were imaged using an epifluorescence microscope (Nikon Eclipse TE2000-5) coupled to a CCD camera (Nikon DS-Qi2) with 20X objective. Fluorescence intensity was quantified using a simple Macro that runs on Image J FIJI software. With this Macro, the background was initially subtracted, and then a region of interest (ROI) was selected (i.e. in the posterior region of the intestine). The mean fluorescence intensity of this selected region was measured. The instructions for creating this macro are available at OSF (https://osf.io/wfgvs/).

### Pharyngeal pumping rate

Feeding rate was quantified essentially as previously described [24]. Briefly, young adult worms (one day after L4 stage synchronization), were transferred to NGM plates seeded with *E. coli* OP50. Animals were filmed under a stereomicroscope at 75× magnification using an Allied Vision Alvium 1800 U-500m camera, at 30 frames per second. Pharyngeal contractions in the terminal bulb were manually counted for 1 minute by analyzing videos at 0.5× playback speed. All experiments were conducted blind to the experimental condition. At least three independent trials with ∼30 animals for each condition were performed.

### Egg Laying Assay

Egg laying behavior was assessed using standard procedures [25]. Briefly, worms were synchronized at the L4 stage. One day after synchronization (young adult stage), individual animals were transferred to NGM plates seeded with *E. coli* OP50 as a food source. After one hour, gravid adults were removed, and the number of deposited eggs was manually counted. At least three independent biological replicates were performed, with 22 worms assessed per experiment.

### Velocity Quantification

Locomotion velocity was quantified using the Multi-Worm Tracker (MWT) (Rex Kerr, https://sourceforge.net/projects/mwt/). Raw tracking data were processed with Choreography (MWT’s feature extraction software) and analyzed using custom MATLAB scripts (The MathWorks, Inc.). To ensure robust tracking, we excluded objects that were not tracked for at least 20 seconds or that moved less than 5 body lengths during the recording.

Animals were placed on bacterial (*E. coli* OP50) lawn-seeded plates (arranged in a donut-shaped pattern to confine food to the edges). A droplet (5ul) of M9 buffer containing the worms was deposited at the center of the plate and allowed to dry before recording. Locomotion was captured at 30 frames per second using an Allied Vision Technology Guppy Pro GPF 125C IRF Camera.

To assess the food-induced slowing response, we computed the ratio between the speed upon encountering the bacterial lawn and the initial speed (movement in the absence of food). A lower ratio indicates a stronger slowing response.

Initial Speed Ratio (%) = (Speed on Food/Initial Speed)×100

## Results

To investigate the neuronal consequences of cytoplasmic TDP-43 accumulation *in vivo*, we generated transgenic *C. elegans* strains expressing human wild-type TDP-43 (*hTDP-43-WT*) or a nuclear localization-deficient variant (*hTDP-43-ΔNLS*) specifically in serotonergic neurons. The serotonergic system in *C. elegans* serves as an ideal platform for investigating neuronal dysfunction, as its activity states are tightly coupled to food availability and directly modulate three core, quantifiable behaviors: (1) feeding rate, (2) locomotion speed, and (3) egg-laying frequency. These stereotypic responses provide a behavioral triad that is highly reproducible across experiments, mechanistically well-characterized, and easily quantifiable through standardized assays [17-22].

In *C. elegans*, feeding rate can be quantified by measuring pharyngeal pumping - the rhythmic contraction of the muscular pharynx, which functions as the worm’s feeding organ [26]. This tubular structure connects the mouth to the intestine and exhibits stereotyped pumping motions that are easily countable under microscopy. The pumping frequency directly correlates with food intake and is modulated by serotonergic signaling [20]. We, therefore, quantified the pharyngeal pumping of young animals on food. We observed striking differences between strains. Wild-type animals displayed robust pharyngeal pumping (333.4 ± 13.8 pumps/min) when exposed to bacterial food (Fig. 1A). As previously reported [27], *tph-1* mutants deficient in serotonin synthesis showed severely impaired pumping (158.2 ± 14.1 contractions/min, p < 0.001 versus wild-type) (Fig. 1A). Animals expressing *hTDP-43-WT* exhibited an intermediate phenotype (283.3 ± 14.1pumps/min), suggesting partial disruption of serotonergic signaling. Most notably, *hTDP-43-ΔNLS* animals showed significantly reduced pumping rates (221.567 ± 12.751 pumps/min, p < 0.001 versus *hTDP-43-WT*) that approached but did not reach the severe deficit observed in *tph-1* null mutants (Fig. 1A).

**Figure 1.**
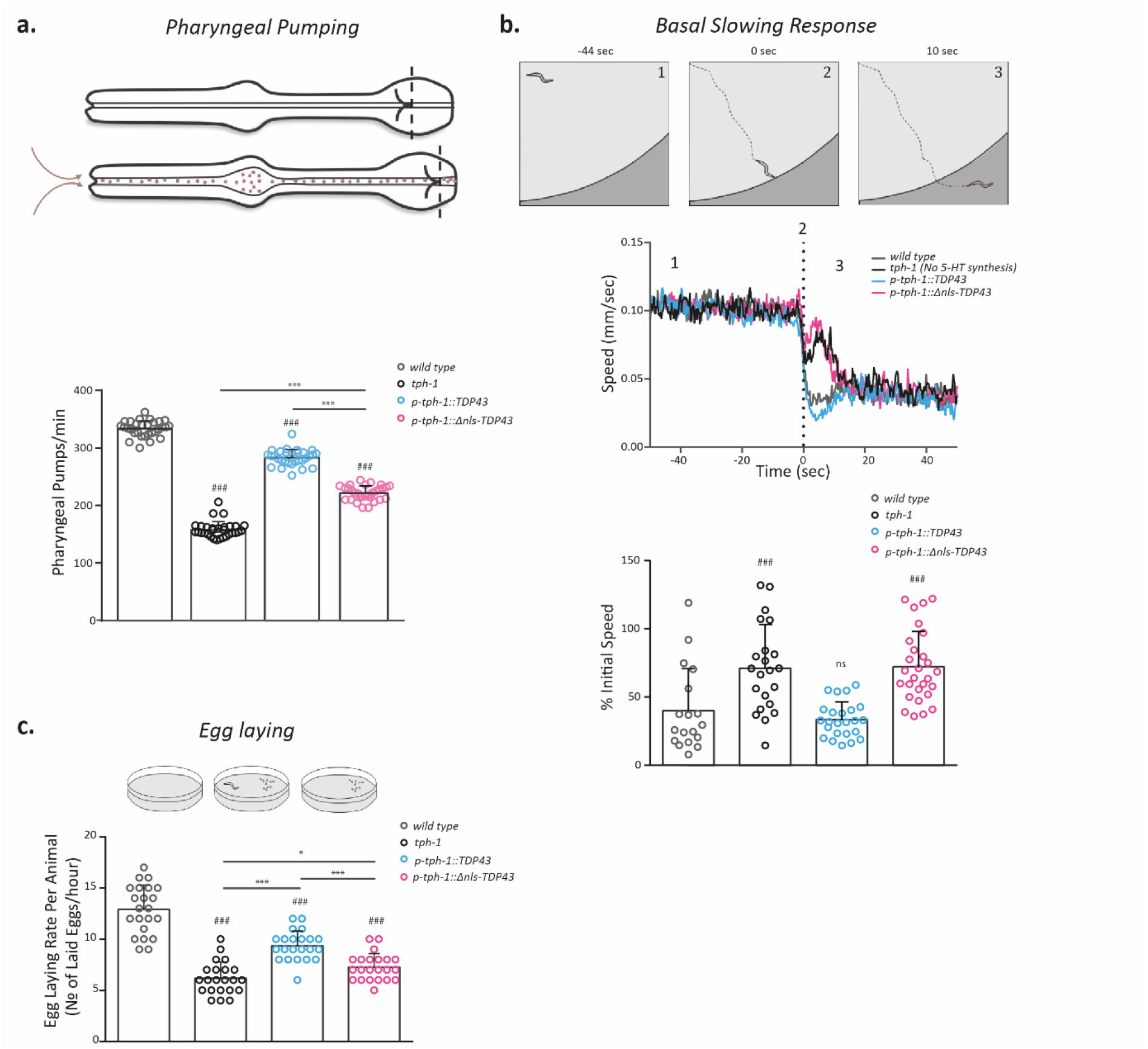
Behavioral effects of nuclear and cytoplasmic *hTDP-43* expression on serotonergic neurons in *C. elegans*. To assess the effect of nuclear, wild-type human TDP-43 (*hTDP-43-WT*) or a cytoplasmic form with a mutated nuclear localization signal (*hTDP-43-ΔNLS*) on serotonergic neuron function in *C. elegans*, we expressed both forms in specific serotonergic neurons and evaluated their impact on feeding and reproductive behaviors. (A) Pharyngeal pumping rate (pumps per minute in the presence of food), a measure of food intake. Left: Schematic of *C. elegans* pharynx. Right: Quantification of pumping rate. n = 30 animals per condition from 3 independent experiments (One-Way ANOVA with Dunn’s post-hoc test). (B) Food-induced slowing response. After 1 hour of food deprivation, speed was measured upon food re-encounter using Worm-Tracker software for video acquisition and MATLAB for analysis. Slowing response was calculated as a percentage of initial speed. n = 26-34 animals per condition from 4 independent experiments (One-Way ANOVA with Dunn’s post-hoc test). (C) Egg-laying frequency. Age-synchronized young adults were transferred to food-seeded plates for 1 hour, followed by egg quantification. n = 22 animals per condition from 3 independent experiments (One-Way ANOVA with Tukey’s post-hoc test). All data is presented as mean ± SD. ^*^p < 0.05, ^**^p < 0.01, ^***^p < 0.001 (comparisons between strains); #p < 0.05 (comparisons versus wild-type).

The enhanced slowing response, where fasted animals abruptly decelerate upon encountering food, is a well-established serotonin-mediated behavior in *C*.*elegans* [17, 18, 28, 29]. To test whether this response was affected by TDP-43 expression, we analyzed locomotion of 1 hour-fasted animals as they approached and encountered bacterial lawns (Fig. 1B). Consistent with previous reports, wild-type animals exhibited a sharp deceleration when approaching the food edge (40.1 ± 30.6% of the initial speed) (Fig. 1B). In contrast, serotonin-deficient *tph-1* mutants exhibited a markedly reduced slowing response (71.1 ± 31.9 % initial speed). While we observed no significant difference between wild-type and *hTDP-43-WT* animals, those expressing *hTDP-43-ΔNLS* in serotonergic neurons exhibited severely diminished slowing, comparable to that of *tph-1* mutants (72.1 ± 26.1).

To complete our functional characterization of serotonergic dysfunction in TDP-43-expressing animals, we quantified egg-laying frequency, a behavior primarily controlled by the HSN serotonergic neurons [30, 31]. As expected, *tph-1* null mutants exhibited dramatically reduced egg-laying rates (6.2 ± 1.6 eggs/hour) compared to wild-type animals (12.9 ± 2.4 eggs/hour; p <0.001), confirming the essential role of serotonin in this behavior (Fig 1C). Animals expressing *hTDP-43-WT* showed an intermediate phenotype (9.4 ± 1.4 eggs/hour; p <0.001 versus wild-type), while those expressing *hTDP-43-ΔNLS* displayed more severe impairment in egg laying (7.3 ± 1.3 eggs/hour; p < 0.001 versus *hTDP-43-WT*). Importantly, the egg-laying defect in ΔNLS-expressing animals, though substantial, remained less severe than in *tph-1* mutants (p= 0.040), suggesting partial preservation of serotonergic signaling (Fig 1C).

Collectively, our behavioral analyses reveal a consistent pattern across three distinct serotonin-dependent processes: feeding, locomotion modulation, and egg-laying. In general, we observed a graded phenotypic severity, with *hTDP-43-ΔNLS* causing more pronounced dysfunction than *hTDP-43-WT*, yet neither variant completely abolished serotonin signaling to the extent seen in *tph-1* null mutants. This spectrum of effects suggests that cytoplasmic accumulation of TDP-43 disrupts serotonergic neuron function, likely through mechanisms that impair but do not fully eliminate serotonin synthesis, release, or reception. The partial retention of function in all assays implies that the observed behavioral deficits reflect specific synaptic or secretory impairments rather than complete neuronal degeneration. These findings position cytoplasmic TDP-43 mislocalization as a key modifier of neuronal dysfunction in proteinopathic states.

To determine whether neurons expressing TDP-43 variants retain functionality, we pharmacologically challenged the animals with fluoxetine and assessed the egg-laying behavior. In wild-type animals, the HSN serotonergic neurons control egg-laying through the coordinated release of serotonin, acetylcholine, and neuropeptides [30, 31]. Fluoxetine is a selective serotonin reuptake inhibitor that robustly stimulates egg-laying [32-34]. Strikingly, this response to fluoxetine only partially depends on serotonin, as *tph-1* mutants (lacking serotonin synthesis) retain significant fluoxetine sensitivity [33]. In contrast, it has been demonstrated that fluoxetine resistance occurred in *egl-1* mutants where HSN neurons are absent, indicating that these neurons mediate both serotonin-dependent and independent components of the response [30, 34, 35]. Unlike *egl-1* mutants, which completely lacked fluoxetine responsiveness, we found that all TDP-43-expressing strains retained some capacity to increase egg-laying following fluoxetine treatment (Fig. 2), indicating that HSN neurons expressing TDP-43 variants maintain basic viability and function. However, quantitative analysis revealed significant differences in the magnitude of this response. While wild-type animals showed robust induction of egg-laying by fluoxetine (11.6 ± 2.0 eggs/hour), *hTDP-43-WT* expressing worms exhibited a markedly reduced response (5.8 ± 1.5 eggs/hour, p<0.01 vs wild-type). Importantly, *hTDP-43-ΔNLS* animals showed even weaker induction (4.5 ± 1.2 eggs/hour, p<0.01 vs *hTDP-43-WT*; p<0.001 vs wild-type), indicating more severe functional impairment. This progressive deficit indicates that cytoplasmic TDP-43 accumulation is a key driver of neuronal dysfunction, even in the absence of complete loss of function.

**Figure 2.**
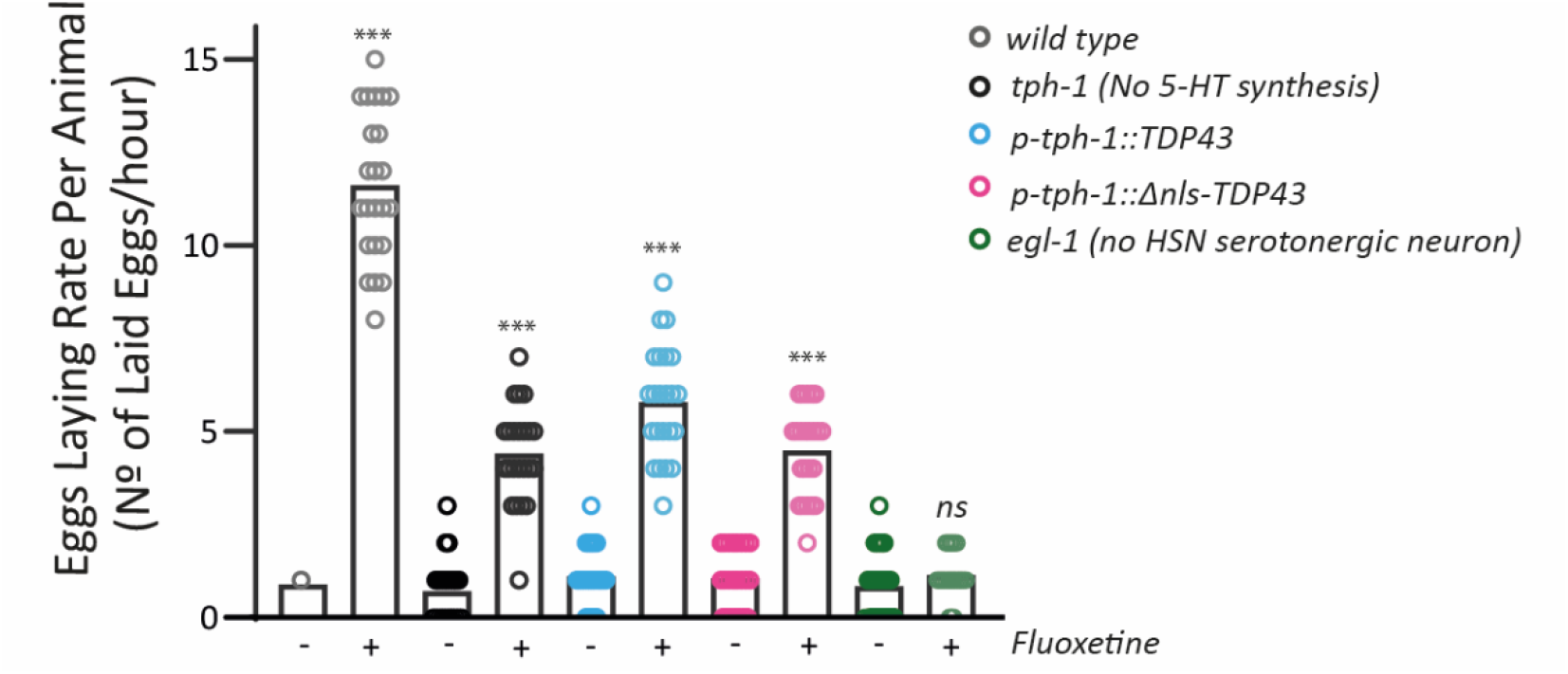
Fluoxetine-induced egg laying in animals expressing pathogenic TDP43 variants in serotonergic neurons. The egg-laying response to fluoxetine (1.6 mM) was evaluated. Fluoxetine induces egg-laying in *C*.*elegans*, and part of its efficacy is dependent on endogenous serotonin (5-HT), correlating with the activity of the serotonin transporter (SERT). The *egl-1* mutant strain, which lacks the serotonergic HSN neuron responsible for egg-laying, was included in the analysis as a negative control. The number of eggs laid within one hour in M9 buffer was quantified. A total of n = 23 animals were analyzed for each condition. A t-test was conducted to compare the egg-laying response in the presence of fluoxetine versus the control condition without fluoxetine. ^*^ shows comparisons between the same strain treated with and without fluoxetine (^***^p<0.001, ^**^p<0.01, ^*^p<0.05; t-test).

Finally, to determine whether the observed functional impairments were associated with structural alterations in serotonergic neurons, we examined neuronal morphology in young adult animals (the same age at which behavioral deficits were detected) using a transcriptional reporter (*Ptph-1::GFP*). This marker selectively labels the serotonergic neurons ADF, NSM, and HSN, enabling detailed visualization of their soma positioning, neurite projections, and overall integrity. Strikingly, we detected no significant morphological abnormalities in any of the examined neurons across strains (Fig. 3 and 4). The somas of ADF, NSM, and HSN neurons were present in 100% of animals, with no differences in their positioning or survival compared to wild-type controls. Furthermore, quantitative analysis of GFP fluorescence intensity—a proxy for neuronal health— revealed no differences between control and TDP-43-expressing strains (^*^p > 0.05 for all comparisons) (Fig. 3 and 4).

**Figure 3.**
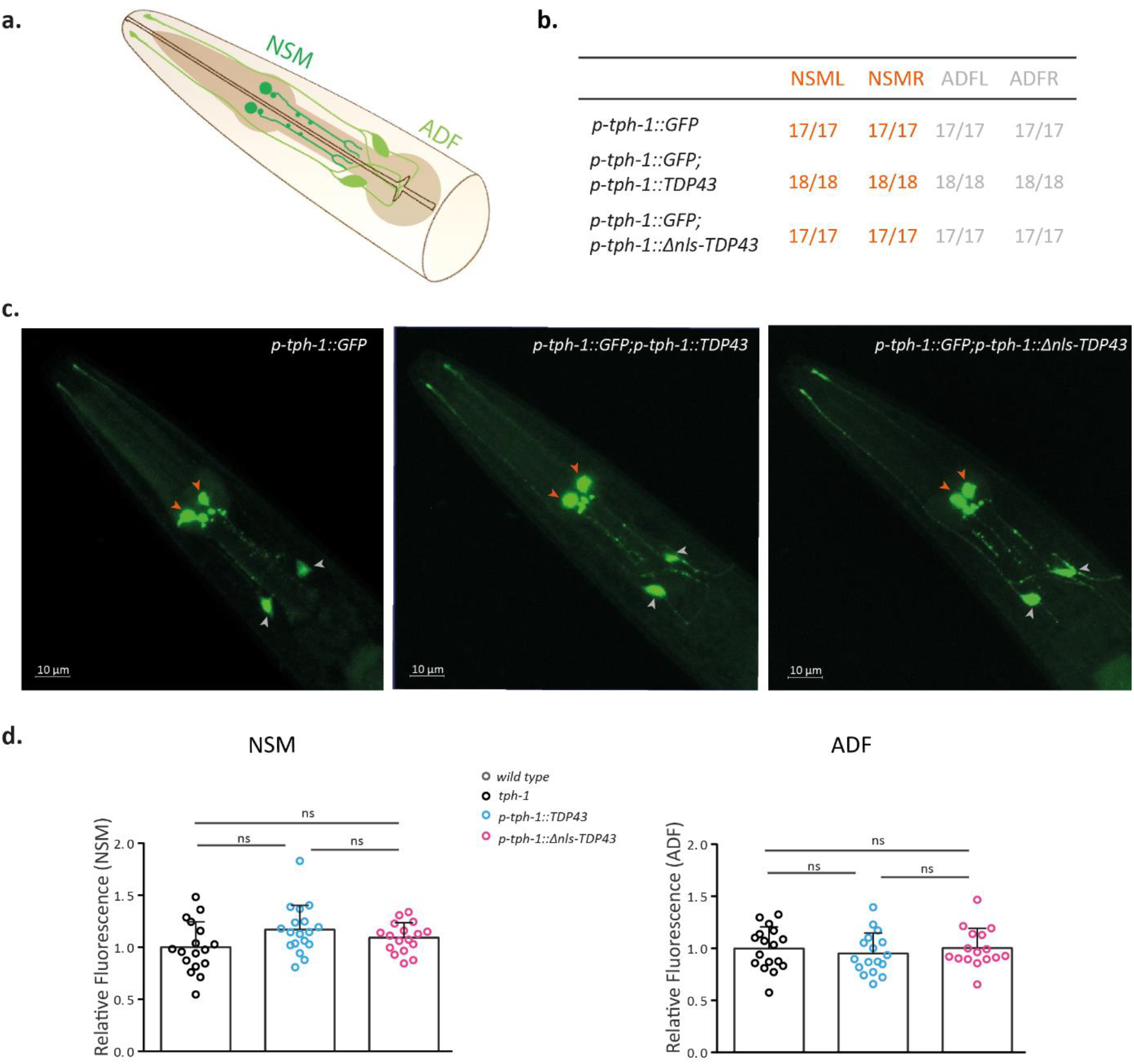
ADF and NSM serotonergic neuron integrity in TDP-43-expressing strains. (A) Schematic of head serotonergic neurons (ADF and NSM) in adult *C. elegans*. (B) Quantification of left and right side neuronal occurrence (worms with detectable neurons/total worms). (C) Representative confocal images showing animals expressing *tph-1*::GFP reporter constructs in wild-type, *hTDP-43-WT* and *hTDP-43-ΔNLS* expressing animals. Neuronal positions are indicated by arrowheads (ADF: white; NSM: orange). Scale bar: 20 μm. (D) Fluorescence intensity quantification of neuronal somata (ImageJ). n = 17-18 animals/condition. One-way ANOVA with Tukey’s test (ns = not significant).

**Figure 4.**
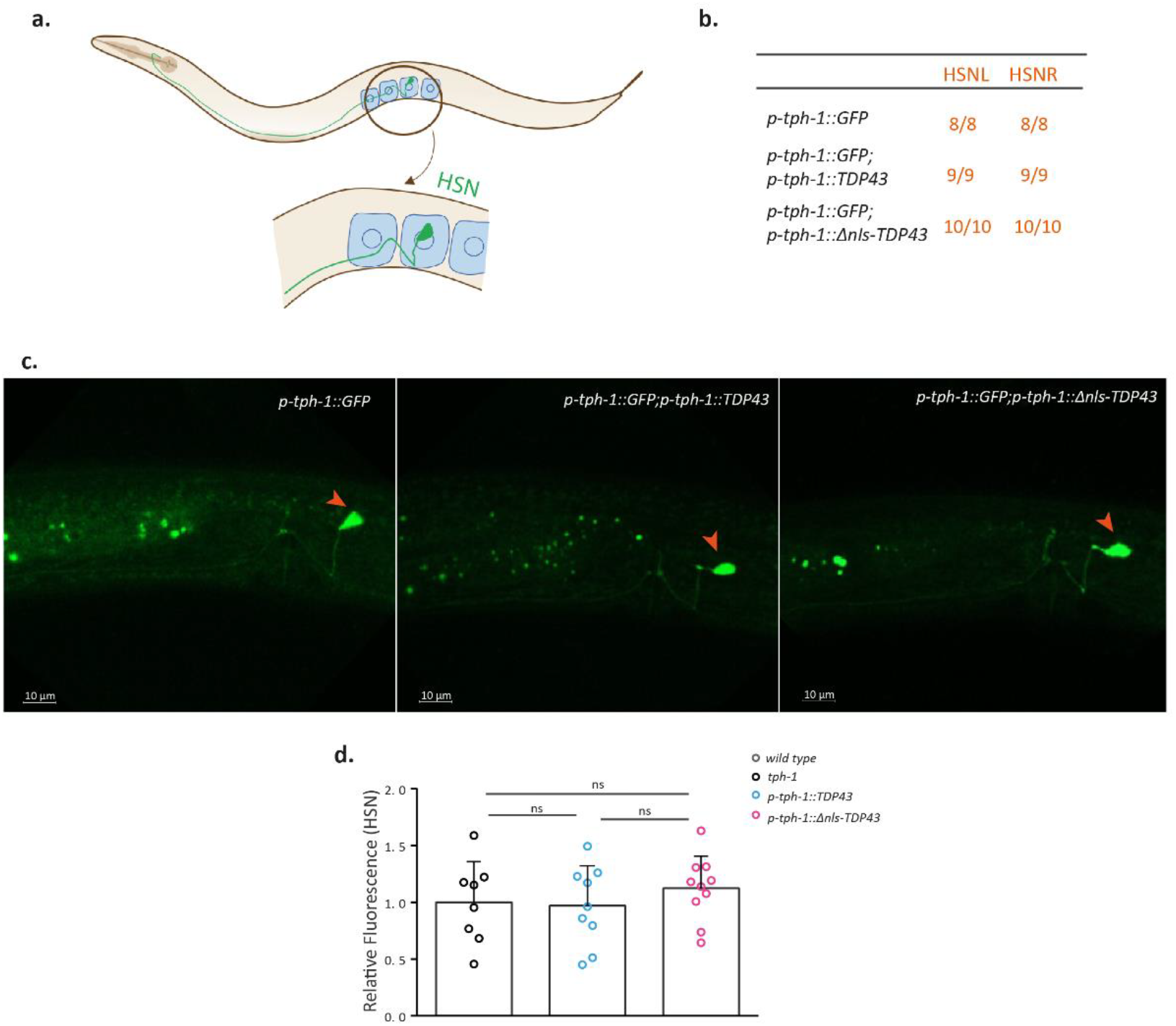
HSN serotonergic neuron integrity in TDP-43-expressing strains. (A) Schematic of the HSN serotonergic neuron position near the vulva in adult *C. elegans*. (B) Quantification of left and right side HSN neuron occurrence (worms with detectable HSN neurons/total worms). (C) Representative confocal images showing HSN neurons (orange arrowheads) in wild-type, *hTDP-43-WT* and *hTDP-43-ΔNLS* expressing animals. (D) Fluorescence intensity quantification of HSN neuronal somata (ImageJ). n = 8-10 animals/condition. One-way ANOVA with Tukey’s test (ns = not significant).

These results demonstrate that the functional deficits caused by *hTDP-43-WT* and *hTDP-43-ΔNLS* expression (e.g., impaired feeding, locomotion modulation, and egg-laying) occur earlier than neuronal loss or gross morphological disruption. The dissociation between behavioral dysfunction and preserved neuronal structure suggests that cytoplasmic TDP-43 accumulation impairs subtle, functional aspects of serotonergic signaling—such as neurotransmitter synthesis, vesicular release, or postsynaptic responsiveness—even before morphological evidence of neurodegeneration becomes apparent.

## Discussion

TDP-43 proteinopathies are a hallmark of several neurodegenerative disorders, including ALS FTD). While traditionally associated with motor neuron degeneration, recent studies have highlighted that other neuronal populations—such as serotonergic neurons—may also be affected in these conditions. For example, reduced serotonergic innervation has been reported in the spinal cord of ALS patients and the brainstem of SOD1(G86R) mouse models [36]. Furthermore, increased expression of 5-HT2B serotonin receptors has been documented in the spinal cord of ALS mice at symptom onset, and Htr2b gene knockout exacerbated disease severity in SOD1(G86R) mice [37, 38].

In this study, we investigated how TDP-43 accumulation affects serotonergic neuron function by expressing either wild-type human TDP-43 (*hTDP-43-WT*) or a nuclear transport-deficient mutant (*hTDP-43-ΔNLS*), which is accumulated in the cytoplasm due to loss of its nuclear localization signal. This targeted approach allowed us to dissect the behavioral and cellular consequences of TDP-43 mislocalization in well-defined neuronal circuits that underlie recognized behaviors in worms.

Our behavioral analyses revealed that both hTDP-43 variants impaired serotonin-dependent behaviors such as pharyngeal pumping, egg-laying, and food-induced locomotion modulation. Notably, expression of cytoplasmic *hTDP-43-ΔNLS* caused more pronounced deficits than *hTDP-43-WT*, with phenotypes that partially mimicked those of *tph-1* mutants, which are unable to synthesize serotonin [27]. However, the impairment of serotonergic signaling is usually milder than that observed in *tph-1* mutants. This graded severity suggests that cytoplasmic mislocalization of TDP-43 compromises serotonergic function, potentially by partially disrupting neurotransmitter synthesis, release, or downstream signaling [39, 40], rather than by inducing cell loss.

Consistent with this hypothesis, we found that *TDP-43*-expressing neurons remained structurally intact. Fluorescence microscopy using a *tph-1*::GFP reporter revealed no overt abnormalities in cell body positioning, axonal projections, or GFP intensity. Moreover, pharmacological exposure to fluoxetine, a selective serotonin reuptake inhibitor, induced egg-laying in *TDP-43*-expressing animals, albeit to a lesser extent than in wild-type controls. This partial responsiveness implies that serotonergic neurons retain the capacity to release neurotransmitters, further supporting the notion of an early functional, rather than structural, impairment. Interestingly, although fluoxetine can activate egg-laying independently of serotonin in *tph-1* mutants, its effects are abolished in *egl-1* animals, which lack the serotonergic HSN neurons altogether [30, 34]. The preserved fluoxetine responsiveness in our TDP-43 models suggests that HSN neurons remain viable and responsive, even in the presence of cytoplasmic TDP-43 accumulation.

Altogether, our results support a model in which cytoplasmic TDP-43 accumulation disrupts serotonergic function without inducing overt neurodegeneration. This dissociation between behavioral dysfunction and structural preservation mirrors clinical findings in early-stage neurodegenerative disease, where functional impairments often precede detectable neuronal loss [41]. Our model thus provides a tractable platform for dissecting the early, circuit-specific effects of TDP-43 pathology and highlights the importance of non-motor neuronal systems in the broader landscape of ALS/FTD pathogenesis.

## Acknowledgments

Some strains were provided by the CGC, which is funded by the NIH Office of Research Infrastructure Programs (P40 OD010440). We thank Maria Jose De Rosa for helpful discussions. In addition, we would like to acknowledge Ignacio Bergé, Edgardo Buzzi, Adrian Bizet, Carolina Gomila, Marta Stulhdreher, María José Tiecher and Carla Chrestía for technical support.

## Funding

This work was supported by Grants from: 1) Agencia Nacional de Promoción de la Ciencia y la Tecnología ANPCYT Argentina to LMI (PICT 2019-1585) and DR (PICT 2019-0480 and PICT-2021-I-A-00052)2) Consejo Nacional de Investigaciones Científicas y Técnicas, Argentina to DR (PIP No. 11220200101606CO) and, 3) Universidad Nacional Del Sur to DR (PGI: 24/B291). The funders had no role in the study design, data collection, and analysis, decision to publish, or preparation of the manuscript.

## Conflict of Interest

The authors declare that the research was conducted in the absence of any commercial or financial relationships that could be construed as a potential conflict of interest.

## Notes

https://osf.io/zpw43/?view_only=9dc9afe51f6a4e3a959d7f17c6f8a8f7

### Competing Interest Statement

The authors have declared no competing interest.

## References

1. Rummens, J. and S. Da Cruz, RNA-binding proteins in ALS and FTD: from pathogenic mechanisms to therapeutic insights. Mol Neurodegener, 2025. 20(1): p. 64.

2. Neumann, M., et al., Ubiquitinated TDP-43 in frontotemporal lobar degeneration and amyotrophic lateral sclerosis. Science, 2006. 314(5796): p. 130–3.

3. Keating, S.S., et al., TDP-43 pathology: From noxious assembly to therapeutic removal. Prog Neurobiol, 2022. 211: p. 102229.

4. Ovsepian, S.V., V.B. O’Leary, and S. Martinez, Selective vulnerability of motor neuron types and functional groups to degeneration in amyotrophic lateral sclerosis: review of the neurobiological mechanisms and functional correlates. Brain Struct Funct, 2024. 229(1): p. 1–14.

5. Yang, L., et al., The Serotonergic System and Amyotrophic Lateral Sclerosis: A Review of Current Evidence. Cell Mol Neurobiol, 2023. 43(6): p. 2387–2414.

6. El Oussini, H., et al., Degeneration of serotonin neurons triggers spasticity in amyotrophic lateral sclerosis. Ann Neurol, 2017. 82(3): p. 444–456.

7. Lu, J., et al., Desloratadine alleviates ALS-like pathology in hSOD1(G93A) mice via targeting 5HTR(2A) on activated spinal astrocytes. Acta Pharmacol Sin, 2024. 45(5): p. 926–944.

8. Altun, Z.F., et al., WormAtlas.

9. Hobert, O., Neurogenesis in the nematode Caenorhabditis elegans. WormBook, 2010: p. 1–24.

10. Pirri, J.K., et al., A tyramine-gated chloride channel coordinates distinct motor programs of a Caenorhabditis elegans escape response. Neuron, 2009. 62(4): p. 526–538.

11. Pirri, J.K., D. Rayes, and M.J. Alkema, A Change in the Ion Selectivity of Ligand-Gated Ion Channels Provides a Mechanism to Switch Behavior. PLoS. Biol, 2015. 13(9): p. e1002238.

12. Chalasani, S.H., et al., Dissecting a circuit for olfactory behaviour in Caenorhabditis elegans. Nature, 2007. 450(7166): p. 63–70.

13. Rani, N., et al., Caenorhabditis elegans: A transgenic model for studying age-associated neurodegenerative diseases. Ageing Res Rev, 2023. 91: p. 102036.

14. Romussi, S., et al., C. elegans: a prominent platform for modeling and drug screening in neurological disorders. Expert Opin Drug Discov, 2024: p. 1–21.

15. Rodriguez, P. and R.D. Blakely, Sink or swim: Does a worm paralysis phenotype hold clues to neurodegenerative disease? J Cell Physiol, 2024. 239(6): p. e31125.

16. Giunti, S., et al., Drug discovery: Insights from the invertebrate Caenorhabditis elegans. Pharmacol Res Perspect, 2021. 9(2): p. e00721.

17. Flavell, S.W., et al., Serotonin and the neuropeptide PDF initiate and extend opposing behavioral states in C. elegans. Cell, 2013. 154(5): p. 1023–1035.

18. Sawin, E.R., R. Ranganathan, and H.R. Horvitz, C. elegans locomotory rate is modulated by the environment through a dopaminergic pathway and by experience through a serotonergic pathway. Neuron, 2000. 26(3): p. 619–631.

19. Churgin, M.A., et al., Antagonistic Serotonergic and Octopaminergic Neural Circuits Mediate Food-Dependent Locomotory Behavior in Caenorhabditis elegans. J Neurosci, 2017. 37(33): p. 7811–7823.

20. Avery, L. and H.R. Horvitz, Effects of starvation and neuroactive drugs on feeding in Caenorhabditis elegans. J. Exp. Zool, 1990. 253(3): p. 263–270.

21. Horvitz, H.R., et al., Serotonin and octopamine in the nematode Caenorhabditis elegans. Science, 1982. 216(4549): p. 1012–1014.

22. Collins, K.M., et al., Activity of the C. elegans egg-laying behavior circuit is controlled by competing activation and feedback inhibition. Elife, 2016. 5.

23. Winton, M.J., et al., Disturbance of nuclear and cytoplasmic TAR DNA-binding protein (TDP-43) induces disease-like redistribution, sequestration, and aggregate formation. J Biol Chem, 2008. 283(19): p. 13302–9.

24. De Rosa, M.J., et al., The flight response impairs cytoprotective mechanisms by activating the insulin pathway. Nature, 2019. 573(7772): p. 135–138.

25. Blanco, M.G., et al., Diisopropylphenyl-imidazole (DII): A new compound that exerts anthelmintic activity through novel molecular mechanisms. PLoS Negl Trop Dis, 2018. 12(12): p. e0007021.

26. Albertson, D.G. and J.N. Thomson, The pharynx of Caenorhabditis elegans. Philos Trans R Soc Lond B Biol Sci, 1976. 275(938): p. 299–325.

27. Sze, J.Y., et al., Food and metabolic signalling defects in a Caenorhabditis elegans serotonin-synthesis mutant. Nature, 2000. 403(6769): p. 560–4.

28. Rhoades, J.L., et al., ASICs Mediate Food Responses in an Enteric Serotonergic Neuron that Controls Foraging Behaviors. Cell, 2019. 176(1-2): p. 85–97 e14.

29. Iwanir, S., et al., Serotonin promotes exploitation in complex environments by accelerating decision-making. BMC Biol, 2016. 14: p. 9.

30. Trent, C., N. Tsuing, and H.R. Horvitz, Egg-laying defective mutants of the nematode Caenorhabditis elegans. Genetics, 1983. 104(4): p. 619–647.

31. Desai, C., et al., A genetic pathway for the development of the Caenorhabditis elegans HSN motor neurons. Nature, 1988. 336(6200): p. 638–646.

32. Choy, R.K. and J.H. Thomas, Fluoxetine-resistant mutants in C. elegans define a novel family of transmembrane proteins. Mol Cell, 1999. 4(2): p. 143–52.

33. Ranganathan, R., et al., Mutations in the Caenorhabditis elegans serotonin reuptake transporter MOD-5 reveal serotonin-dependent and -independent activities of fluoxetine. J. Neurosci, 2001. 21(16): p. 5871–5884.

34. Dempsey, C.M., et al., Serotonin (5HT), fluoxetine, imipramine and dopamine target distinct 5HT receptor signaling to modulate Caenorhabditis elegans egg-laying behavior. Genetics, 2005. 169(3): p. 1425–36.

35. Conradt, B. and H.R. Horvitz, The C. elegans protein EGL-1 is required for programmed cell death and interacts with the Bcl-2-like protein CED-9. Cell, 1998. 93(4): p. 519–529.

36. Dentel, C., et al., Degeneration of serotonergic neurons in amyotrophic lateral sclerosis: a link to spasticity. Brain, 2013. 136(Pt 2): p. 483–93.

37. El Oussini, H., et al., Serotonin 2B receptor slows disease progression and prevents degeneration of spinal cord mononuclear phagocytes in amyotrophic lateral sclerosis. Acta Neuropathol, 2016. 131(3): p. 465–80.

38. Arnoux, A., et al., Evaluation of a 5-HT2B receptor agonist in a murine model of amyotrophic lateral sclerosis. Scientific Reports, 2021. 11(1): p. 23582.

39. Lépine, S., et al., Homozygous ALS-linked mutations in TARDBP/TDP-43 lead to hypoactivity and synaptic abnormalities in human iPSC-derived motor neurons. iScience, 2024. 27(3): p. 109166.

40. Dyer, M.S., et al., Mislocalisation of TDP-43 to the cytoplasm causes cortical hyperexcitability and reduced excitatory neurotransmission in the motor cortex. J Neurochem, 2021. 157(4): p. 1300–1315.

41. Gelon, P.A., P.A. Dutchak, and C.F. Sephton, Synaptic dysfunction in ALS and FTD: anatomical and molecular changes provide insights into mechanisms of disease. Front Mol Neurosci, 2022. 15: p. 1000183.

